# Transient resting-state network dynamics in cognitive ageing

**DOI:** 10.1101/2020.05.19.103531

**Authors:** Roni Tibon, Kamen Tsvetanov, Darren Price, David Nesbitt, Cam-CAN, Richard Henson

## Abstract

It is important to maintain cognitive function in old age, yet the neural substrates that support successful cognitive ageing remain unclear. One factor that might be crucial, but has been overlooked due to limitations of previous data and methods, is the ability of brain networks to flexibly reorganise and coordinate over a millisecond time-scale. Magnetoencephalography (MEG) provides such temporal resolution, and can be combined with Hidden Markov Models (HMMs) to characterise transient neural states. We applied HMMs to resting-state MEG data from a large cohort (N=594) of population-based adults (aged 18-88), who also completed a range of cognitive tasks. Using multivariate analysis of neural and cognitive profiles, we found that decreased occurrence of “lower-order” brain networks, coupled with increased occurrence of “higher-order” networks, was associated with both increasing age and impaired fluid intelligence. These results favour theories of age-related reductions in neural efficiency over current theories of age-related functional compensation.

## Main

With the increasing proportion of older adults in the worldwide population^1^, there is a pressing need to understand the neurobiology of cognitive ageing. Normal ageing generally results in cognitive decline^2^, though not all cognitive functions follow the same trajectory. In particular, whereas age produces a marked impairment in fluid intelligence (the ability to solve new problems), crystallised intelligence (the ability to rely on acquired knowledge) shows more modest age-related changes^3–7^. Moreover, functional neuroimaging has revealed that ageing is associated with changes in connectivity between brain regions, both within and between large-scale networks^8^. One factor that might play a crucial role in the ability to maintain cognition in old age, but which has been largely overlooked, is the ability of brain networks to flexibly reorganize and coordinate on a sub-second time-scale. Indeed, the relationship between cognition and such transient brain connectivity, and how this relationship differs with age, remains unknown.

In recent decades, functional connectivity within the human brain has been measured mainly with functional magnetic resonance imaging (fMRI). In particular, differences in the brain’s connectivity during the resting-state (rsfMRI) have proved effective in distinguishing various patient groups from controls (e.g., Alzheimer disease, major depression, schizophrenia; see for example^9^). Substantial work has also used rsfMRI to examine age-related changes in functional connectivity^10–15^. However interpreting such connectivity changes measured with fMRI is difficult owing to methodological challenges, such as confounding factors like vascular reactivity and head motion, which also change with age^16–19^. While some of these confounds, like neurovascular coupling, can be addressed by more sophisticated modelling^20^, the fact remains that fMRI has a fundamentally limited temporal resolution (owing to the sluggish vascular response and relatively slow image acquisition times), which precludes it from disclosing the potentially richer dynamics in brain connectivity above approximately 0.1 Hz.

An alternative, non-invasive way of measuring functional connectivity is offered by magnetoencephalography (MEG), which can sample neural activity at 1kHz and higher (at the cost of worse spatial resolution). Indeed, recent advances in analytical approaches offer the ability to measure *dynamic* functional connectivity, i.e., “microstates” of stable connectivity patterns that last a few hundred milliseconds^21^, well beyond the temporal resolution of fMRI. Importantly, MEG is less sensitive than fMRI to age-related changes in vascular factors and head motion. In the current study, we utilized these advantages of MEG to relate transient resting-state dynamics to cognitive ageing.

More specifically, we exploited a large resting-state MEG (rsMEG) dataset obtained from 594, population-based individuals sampled uniformly across the adult-lifespan (18 to 88 years of age) as part of the Cam-CAN project (www.cam-can.org). In addition to the rsMEG scan, these individuals also completed a wide range of cognitive tasks. We characterised transient network dynamics using the recent application of Hidden Markov Models (HMMs) in order to explore the temporal dynamics of rsMEG networks^21–26^. An HMM is a data-driven method that identifies a sequence of “states”, where each state corresponds to a unique pattern of brain covariance that reoccurs at different points in time. By quantifying the time-series of MEG data as a sequence of transient states, the HMM provides information about the periods of time at which each state is active, enabling the characterisation of its temporal dynamics. While this technique has identified network dynamics in small resting-state or task MEG datasets^21,23,24^, these dynamics have not yet been linked to age and cognition. In particular, the size of the Cam-CAN cohort allowed us to take a multivariate approach, namely to use canonical correlation analysis (CCA) to relate the temporal properties of the data-driven HMM states to profiles of cognitive performance, and to test whether these relations differ with age.

## Results

Results were obtained following the data pre-processing and analyses procedures summarised in Figure 1 below. The raw data can be obtained by applying for access through the Cam-CAN data portal (https://camcan.mrc-cbu.cam.ac.uk/). All code used in the paper will be available via an online repository.

**Figure 1.**
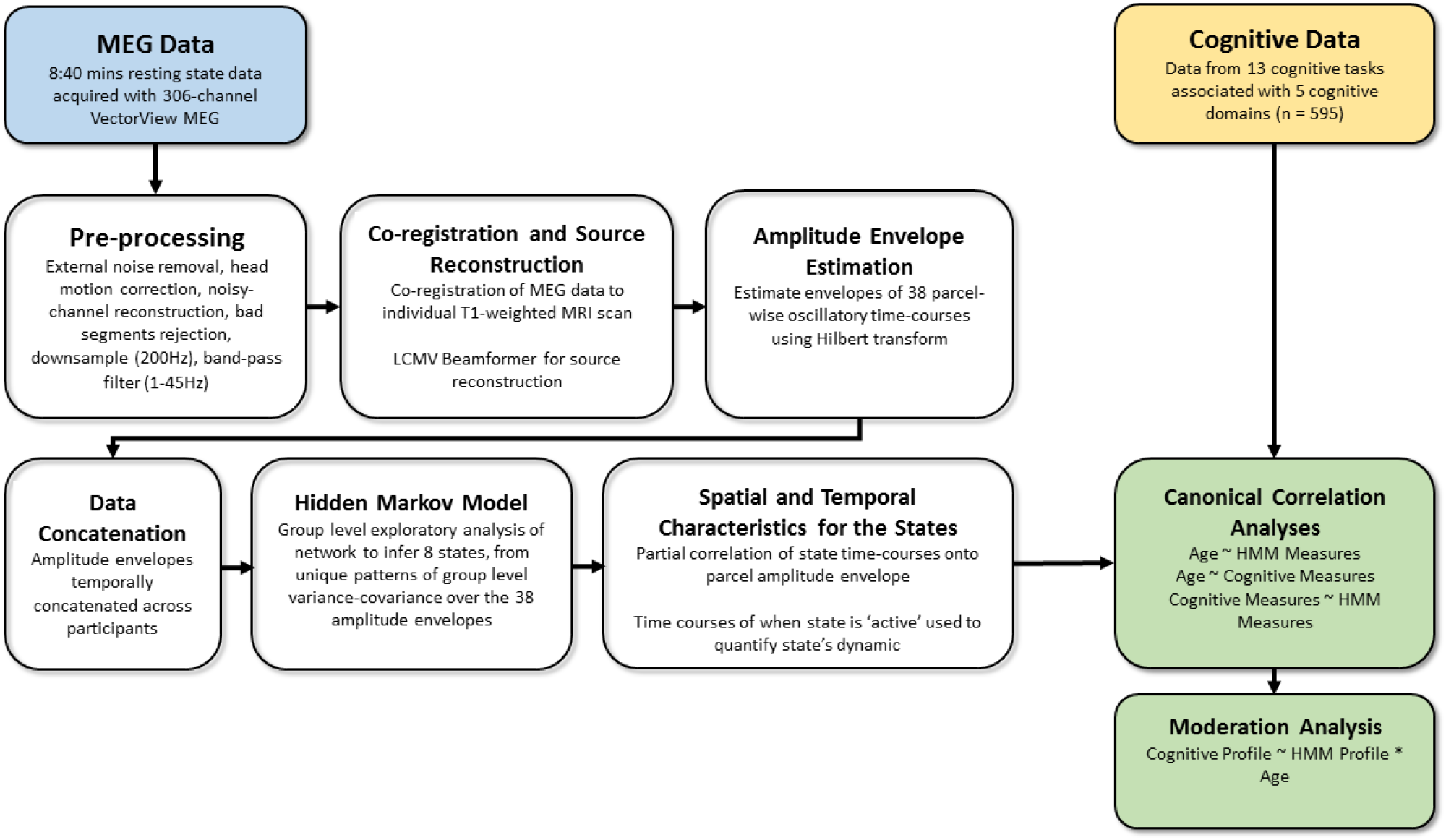
Overview of processing and analysis pipeline used in the study.

Figure 2 shows the spatial maps of the 8 networks (states) derived from combing the MEG data across all participants. The states include three distributed frontotemporoparietal networks (FTP1, FTP2, FTP3), a higher-order visual network (HOV), two early visual networks (EV1, EV2) and two sensorimotor networks (SM1, SM2). They are similar to those obtained from young adults in previous studies^21–23^.

**Figure 2.**
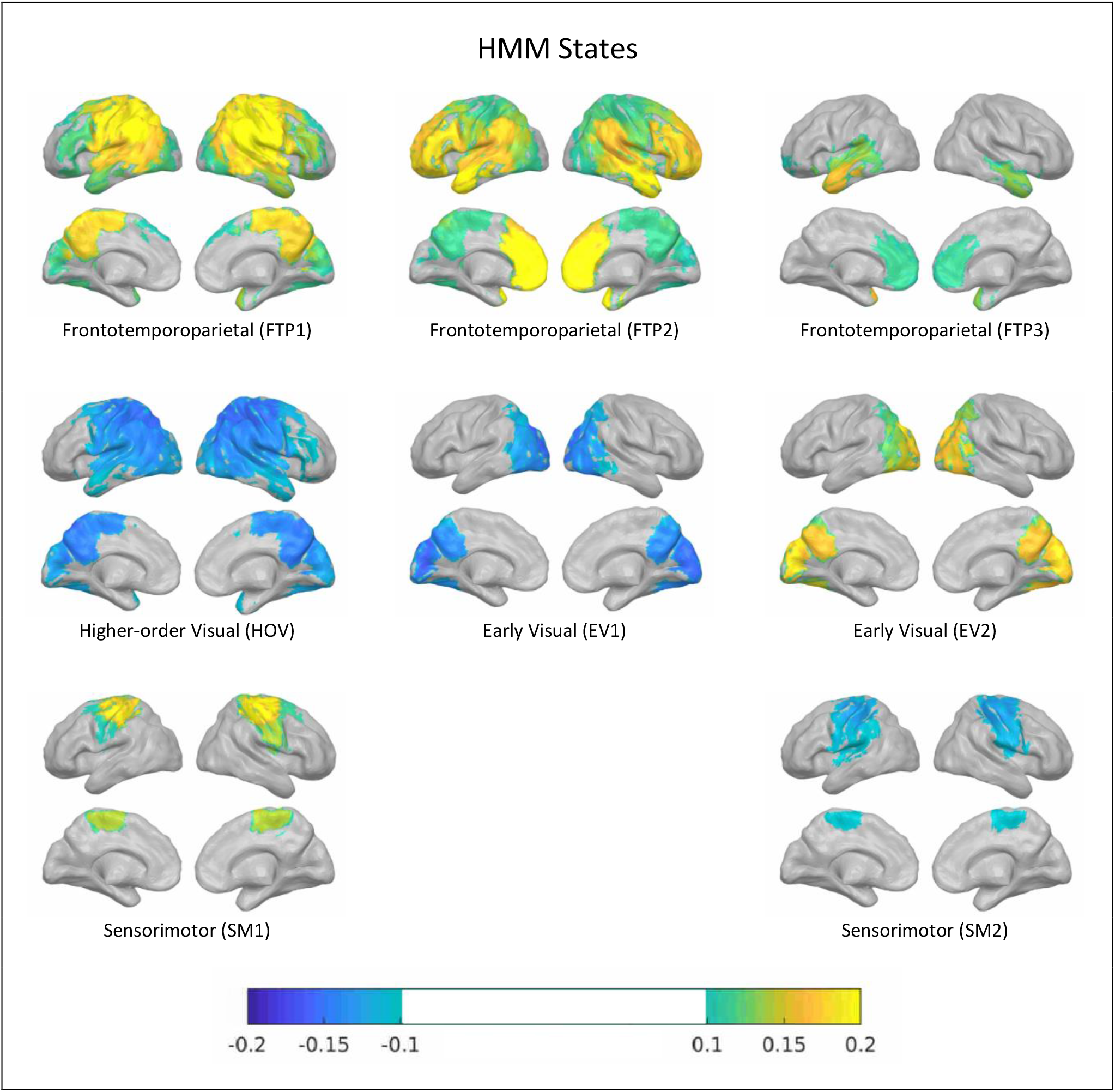
The 8 inferred HMM states. Each map shows the partial correlation between the state time course and the parcel-wise amplitude envelopes. Yellow colours represent amplitude envelope increases when the brain visits that state and blue colours represent envelope decreases. The partial correlation values have been thresholded to show correlation values above 50% of the maximum correlation across all states. To refer to the states, we use the same naming scheme applied by Hawkins et al.^23^.

We next characterized the temporal properties of each state in terms of 4 metrics: fractional occupancy (FO), mean life time (MLT), number of occurrences (NO) and mean interval length (MIL). Group averages for each measure in each state are shown in Figure 3. Overall, primary (visuo-motor) states had higher number of occurrences than higher-order states. The most commonly-occurring networks were sensorimotor network SM2 and early visual network EV1, which had the highest mean FO and NO. The network with the most prolonged visits (highest MLT) was the high-order visual network HOV. Conversely, the least common network, with lowest FO and NO and with greatest MIL, was frontotemporoparietal network FTP3. These findings largely agree with Hawkins et al.^23^ and (to a lesser extent) with other previous studies^21,22^, though now based on a much larger sample with a much larger age range.

**Figure 3.**
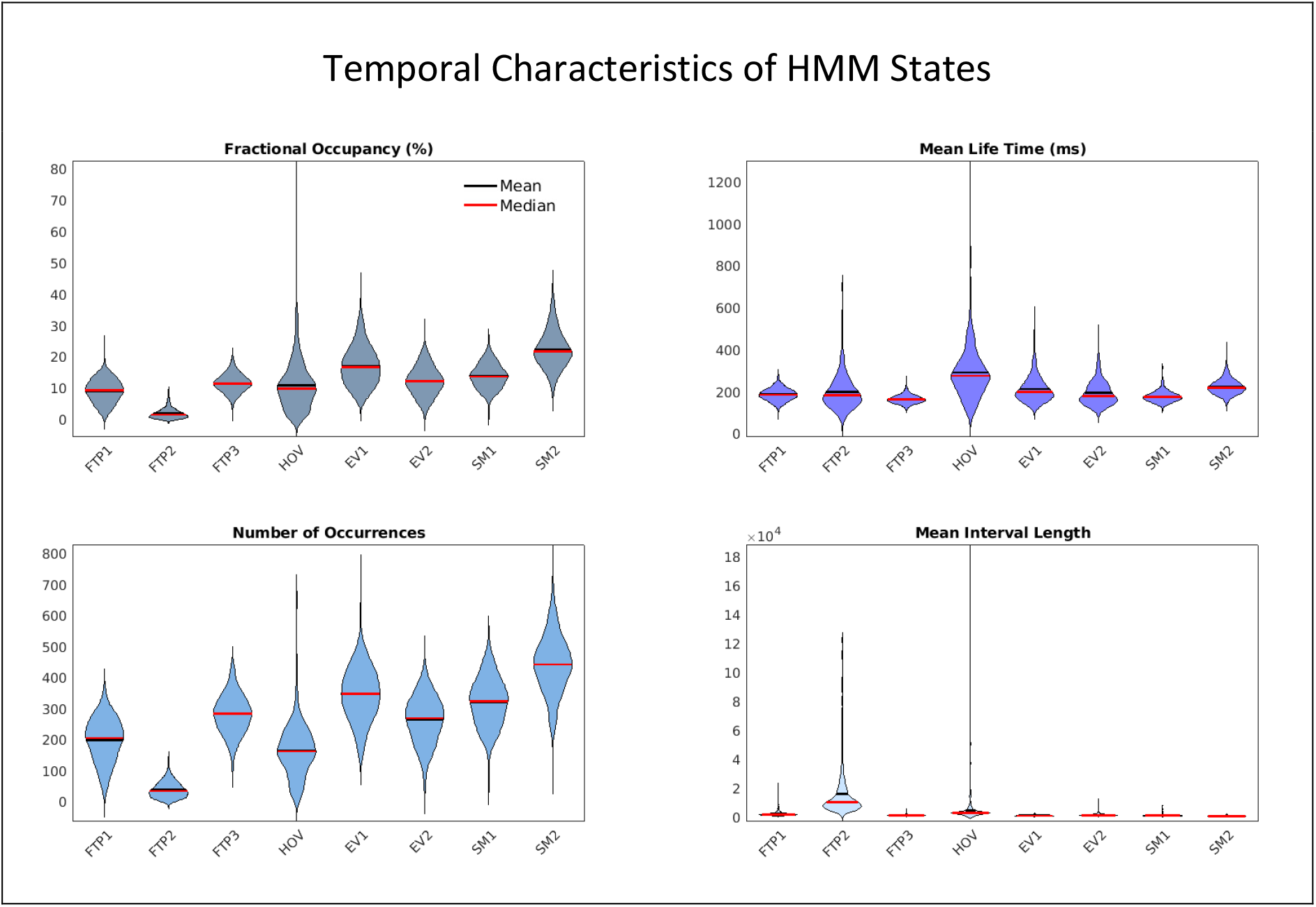
Violin plots^27^ of the four temporal characteristics of the HMM states: % fractional occupancy (FO; top-left), mean life time (MLT; top right), number of occurrences (NO; bottom-left) mean interval length (MIL; bottom-right). The first three measures are positive measures (i.e., indicate more frequent/longer duration of state’s occurrence), whereas the fourth measure (MIL) is a negative measure. Mean and median are indicated by black and red lines, respectively (N=595).

Our next step was to apply CCA to relate the 32 temporal characteristics of the HMM states (4 metrics for each of the 8 states) to age (such a multivariate analysis is important in the presence of multicollinearity, given the co-dependence between the 4 metrics of HMM state dynamics). One participant who had no visits to one of the states (HOV) was excluded from these analyses. Only a single CCA mode was possible (given the unidimensional age variable), which explained 28.6% of the combined variance (*R_c_* = .53, *p* < .001). Table 1 shows the structure coefficients (r_s_) for each metric of each state. As apparent in the table, the three frontotemporoparietal states (FTP1, FTP2, FTP3) and the higher-order visual state (HOV) showed positive correlations with age for FO, MLT and NO measures, and tended to show negative correlation for the MIL measure, whereas the two early visual states (EV1 and EV2) and one sensorimotor state (SM1) tended to show negative correlations with age for FO, MLT, NO and positive correlation for MIL. In other words, older people had more and longer occurrences of states involving frontotemporoparietal and higher-order visual states, and fewer, shorter occurrences of early visual and motor states (with the exception of sensorimotor state SM2, which did not show a significant relationship with age).

**Table 1.** Structure coefficients for the CCA relating HMM measures with age (N=594)

|  |  |  |  |  |  |
| --- | --- | --- | --- | --- | --- |
| State | FO | MLT | NO | MIL | Age |
| FTP1 | .27** | .13* | .28** | -.12* | (1) |
| FTP2 | .23** | .15** | .24** | -.21** |  |
| FTP3 | .2** | .44** | .12* | -.07 |  |
| HOV | .23** | .15** | .34** | -.07 |  |
| EV1 | -.35** | -.5** | -.00 | -.01 |  |
| EV2 | -.45** | -.38** | -.33** | .18** |  |
| SM1 | .18** | .04 | .22** | -.13* |  |
| SM2 | -.03 | -.09* | .00 | .02 |  |

\* $p < .05$ , \*\* $p < .005$ . Note: Each cell depicts structure coefficients ( $r_s$ ). Coefficients are shown for each of the 4 HMM measures (FO, MLT, NO, MIL), for each state. $r_s$ for Age is 1, because this set contains only one variable.

Once we established age effects on the pattern of occurrence of brain states, we asked how the pattern of the states’ occurrence relates to cognition. To this end, we related the temporal characteristics of the states to the 13 cognitive measures (see Methods). 12 CCA modes showed a significant correlation coefficient (Bonferroni corrected p-value across 13 modes < .05). Nevertheless, given the relatively low squared canonical correlation (*R^2^_c_*) of most modes (see Supplementary Figure 1), only the first mode which accounted for a substantial portion of the combined variance (24%) was explored further (see Sherry & Henson^28^ for a similar approach). Figure 4 presents the structure coefficients (*r_s_*) of the first CCA mode. For the cognitive data, the coefficients resembled the pattern associated with poor (low) fluid intelligence reported before^3^. Specifically, *r_s_* was negative for all the fluid intelligence tests except the multitasking (MltTs) and motor speed (MRSp) tests, where higher values of these response-time measures represent worse performance, and the proverbs (ProV) and Spot-the-Word (STW) tests, which measure crystallised intelligence instead. Importantly, the four “higher-order” states (FTPs and HOV) were positively related to this cognitive profile, i.e., greater occurrence of these states (as indicated by increased FO, MLT, NO and decreased MIL) was associated with lower fluid intelligence. Furthermore, this cognitive profile was associated with decreased activation of the two early visual states (EVs). The two sensorimotor states did not show strong relations to this cognitive profile.

Finally, we asked whether the relationship between the brain profile of HMM measures and the cognitive profile differed with age. For this moderation analysis, we constructed a multiple linear regression model that included participants’ scores for the HMM profile, their age and their interaction (HMM profile × age) as predictors, and participants’ scores for the cognitive profile as the dependent variable. The HMM scores were significantly associated with cognitive scores after accounting for the main effect of age (*β* = .13, *t*(590) = 4.51, *p* < .001), demonstrating that the above brain-cognitive relationship was not driven solely by age effects. Moreover, the interaction between age and HMM profile was also significant (*β* = .11, *t*(590) = 4.33, *p* < .001), demonstrating that the brain-cognition relationship was moderated by age in a positive sense, i.e., becoming more positive with age. To visualise this moderation effect, the brain-cognition relation was repeated for six equally-sized age groups (n=99 for each group; 18–34 years old; 34-45 years old; 45-55 years old; 55-66 years old; 66-76 years old; 76-88 years old). As shown in Figure 4, the brain-cognition relationship was stronger in the older groups.

**Figure 4.**
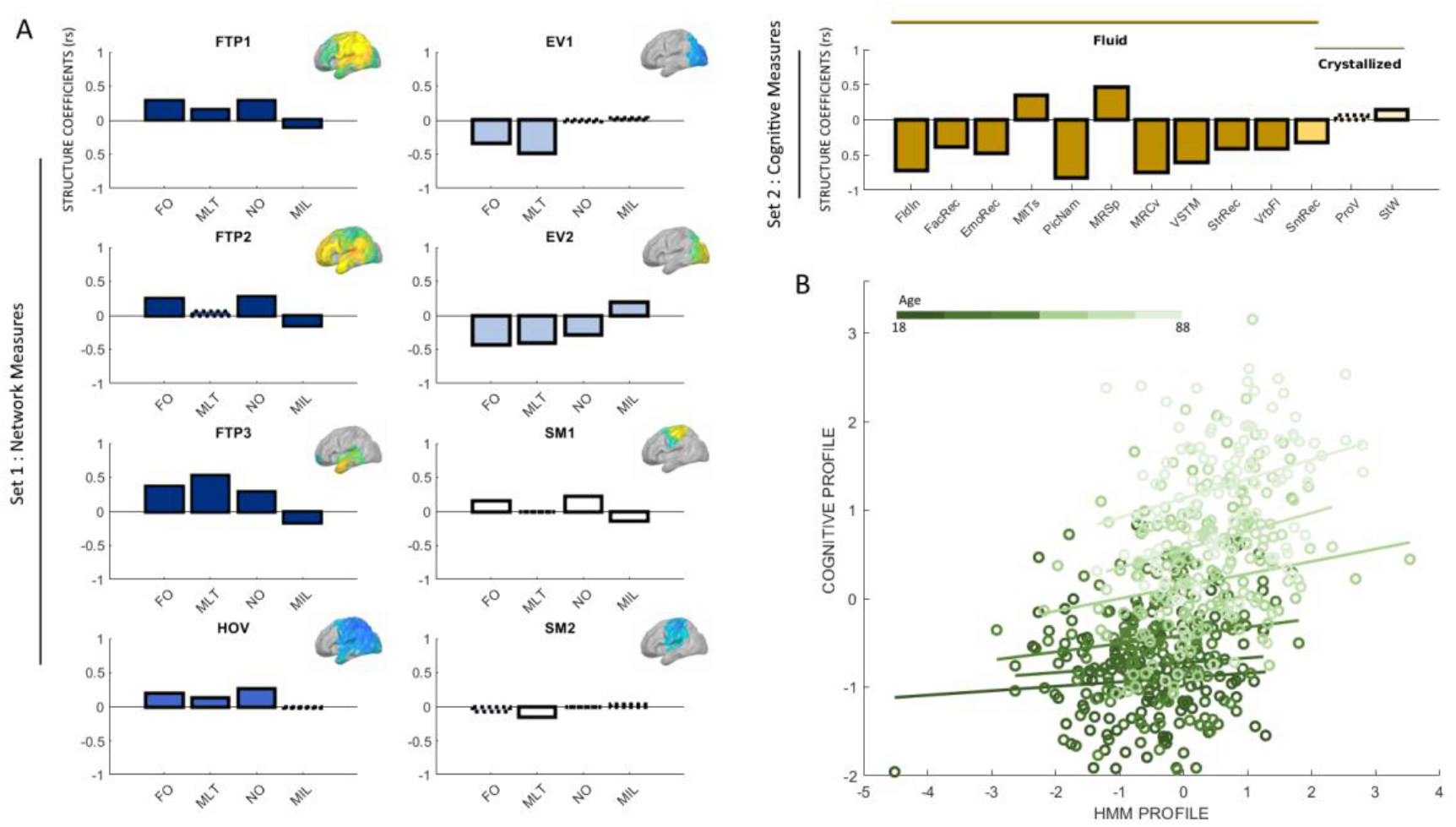
Outcomes of main CCA and moderation analyses (N=594). **(A)** Structure coefficients (*r_s_*) for the CCA relating HMM measures with cognitive measures. Solid outlines represent statistically significant r_s_, whereas dashed outlines represent r_s_ that are not statistically significant. r_s_ for brain HMM measures are shown in blue/white, with different shades of blue representing different types of networks (FTP, HOV etc). Corresponding HMM state maps are inset. For clarity, r_s_ for each network are shown separately, though in practice all were included in a single CCA analysis. r_s_ for the cognitive measures are shown in brown, with different shades indicating the distinction between cognitive abilities obtained via confirmatory factor analysis in Borgeest et al.^3^: fluid intelligence (dark brown), crystallised intelligence (light brown) and mixed (intermediate brown; for SntRec). For the response-time measures of MltTs and MPSp, lower scores indicate better performance (hence the opposite sign). **(B)** Scatter plot of bivariate correlations for six age groups. Dark shades of green represent younger adults, whereas light shades represent older adults. The relationship between HMM and cognitive profiles is higher for older adults (formally confirmed by a continuous moderation analysis; see text).

## Discussion

Our results show that transient neural dynamics, particularly those of high-order and early-visual states, differ across the healthy adult lifespan, with an increasing importance for cognitive function in older than younger adults. Importantly, by using a novel data-driven method to infer brain states from MEG data, we were able to overcome some of the limitations of the more common use of fMRI to examine ageing and cognition, such as confounding effects of vascular health, head motion and the ability to examine only very slow dynamics owing to low-frequency fluctuations of the fMRI response. More specifically, our finding that age and impaired fluid intelligence are associated with increased occurrence of brain states involving “higher-order” networks (such as those straddling frontal, temporal and parietal cortex) are less consistent with theories of functional compensation in ageing, and more consistent with theories of reduced neural efficiency in ageing, as we expand below.

We used multivariate analysis (CCA) to relate the pattern of brain dynamics (HMM profile) to age and to cognition, and to examine whether the brain-cognition relationship was moderated by age. This first analysis relating brain dynamics to age showed that the occurrence (e.g., frequency and duration) of states involving frontotemporoparietal and higher-order visual regions increase with age, whereas the occurrence of states involving early-visual regions decrease with age. The second analysis relating brain dynamics to cognitive performance revealed that a similar profile of increased occurrence of higher-order states and reduced occurrence of early-visual states is associated with a pattern of poorer performance on tests of fluid intelligence, but not tests of crystallised intelligence. Importantly, our final moderation analysis showed that this relationship between brain and cognitive profiles is not simply a result of shared influences of age (since age was included as a covariate). Moreover, this moderation analysis showed that the brain-cognitive relationship gets stronger with increased age, such that reduced cognitive performance in older participants is more strongly associated with the shift from early visual to higher-order networks than in younger participants.

To our knowledge, this is the first study to report a resting state shift from lower to higher-order networks that is linked to both age and cognition. This finding nevertheless shares some similarity with previous fMRI findings. For example, Davis et al.^29^ summarised a pattern across a number of fMRI experiments (first observed by Grady et al.^30^) in which older adults show increased activity in anterior (e.g., frontal) brain regions, and decreased activity in posterior (e.g., visual) brain regions, and called this the “Posterior-to-Anterior Shift with Ageing” (PASA). Furthermore, it has been hypothesised that one reason for this shift is “functional compensation”, whereby older people activate frontal regions to compensate for age-related impairments in posterior brain regions, that is, in order to maintain levels of cognitive performance^29,31,32^. However, this pattern and interpretation are based on activations during task-based fMRI, i.e., while participants are performing a cognitive task, rather than the current resting state. Moreover, more recent fMRI studies have questioned whether PASA reflects functional compensation, and suggest instead that the increased frontal activation reflects reduced neural efficiency or specificity^33–38^. The difference between these two interpretations of the PASA finding is that, whereas the functional compensation hypothesis predicts that anterior increases correlate with better cognitive performance, the inefficiency hypothesis predicts the opposite pattern of anterior increases correlating with worse cognitive performance. Our findings support the latter account, i.e., that increased occurrence of higher-order states is associated with worse cognitive performance, specifically in measures of fluid intelligence and particularly in older adults. Importantly, unlike previous reports of the PASA pattern, the current shift was observed during rest, suggesting that it might reflect a stable property of the ageing brain.

The results of our moderation analysis resemble those obtained in a previous study^39^, which also showed that the relationship between brain connectivity and cognition increased with age, but using rsfMRI instead. In that case, the authors showed that the cognitive function of older adults becomes increasingly dependent on the balance of excitatory connectivity between networks, and the stability of intrinsic neural representations within networks. Importantly, they used biophysical modelling to account for the confounding effects of vascular health on the fMRI response. However, their results were still limited to static connectivity driven by the low-frequency fluctuations that are visible to fMRI, and to a small subset of three brain networks (owing to the complexity of the biophysical modelling). The current study overcomes these issues by utilising rsMEG to measure i) neural activity directly, rather than indirectly via a vascular response, ii) dynamic connectivity with much higher temporal resolution and iii) whole-brain networks (within the limits of MEG resolution).

One caveat of the current study is that we used cross-sectional data, which precludes direct inferences about ageing (as distinct from cohort effects, owing to year of birth). However, we are not aware of any longitudinal MEG data on such a large, representative population, and until such time, our results can be used to justify and inform hypotheses for future rsMEG studies of ageing.

Furthermore, the HMM approach comes with some assumptions. For example, it relies on group concatenation that here assumes anatomical correspondence across the lifespan (though our use of relatively large ROIs minimizes this issue), it requires a priori specification of the number of states, and it uses a Gaussian observation model which may be an oversimplification of the underlying network dynamics^21^. Nevertheless, despite these limitations, our study offers novel insights on the relationship between the cognitive sequelae of ageing and the underlying patterns of functional brain dynamics, which may be used in the future for mechanistic justification and assessment of interventions to reduce the personal and societal burden of cognitive impairments in old age.

## Online Methods

### Participants

A population-based sample of 708 healthy human adults (359 women and 349 men) was recruited as part of Stage 2 of the Cambridge Centre Aging and Neuroscience (Cam-CAN; www.cam-can.org^40^). Ethical approval for the study was obtained from the Cambridgeshire 2 (now East of England-Cambridge Central Research Ethics Committee), and participants gave full informed consent. Exclusion criteria included poor vision (below 20/50 on Snellen test^41^) and poor hearing (threshold 35 dB at 1000 Hz in both ears), ongoing or serious past drug abuse as assessed by the Drug Abuse Screening Test (DAST-20^42^), significant psychiatric disorder (e.g., schizophrenia, bipolar disorder, personality disorder), neurological disease (e.g., known stroke, epilepsy, traumatic brain injury), low score in the Mini Mental State Exam (MMSE; 24 or lower^43^), or poor English knowledge (non-native or non-bilingual English speakers); a detailed description of exclusion criteria can be found in Shafto et al^40^, Table 1. Of these, only participants with full neuroimaging data (resting state MEG data and structural MRI data) were considered for the current study (*n* = 610). Fifteen additional participants were excluded from analyses due to poor MEG-MRI co-registration (details below). Thus, the final sample included 595 participants (299 women and 296 men, age range 18-88).

### Cognitive Tasks

13 cognitive tasks, performed outside the scanner, were used to assess five broad cognitive domains, including executive functions, memory, language functions, processing speed, and emotional processing. The tasks are summarized in Table 2, and are fully detailed in Shafto et al^40^. Task data were obtained from Borgeest et al.^3^, in which missing data (<12% in all tasks) were interpolated using the Full Information Maximum Likelihood (FIML) method^44^ in the Lavaan R package^45^ to allow unbiased estimates, applied to the full Stage 2 sample (*n* = 708).

**Table 2.** Description of cognitive behavioural tasks (table adapted from Borgeest et al.^3^)

| Cognitive Domain | Cognitive Task | Task Description | Descriptive Statistics for $n=595$ (Mean, SD, Range) | References |
| --- | --- | --- | --- | --- |
| Executive Function | Fluid Intelligence (FldIn) | Cattell Culture Fair Test, incl. nonverbal puzzles involving series completion, classification, matrices, and conditions. | $M = 31.74$<br>$SD = 6.83$<br>$Range = 11-44$ | Cattell & Cattell, 1960 |
| | Multitasking (Hotel Task; MItTs) | Perform tasks in role of hotel manager: write customer bills, sort money, proofread advert, sort playing cards, alphabetise list of names. Total time must be allocated equally between tasks; there is not enough time to complete any one task. | $M = 301.3$<br>$SD = 171.7$<br>$Range = 20.19-960$ | Shallice & Burgess, 1991 |
| Language Functions | Spot the Word (StW) | Involves presenting an individual with pairs of items comprising one word and one non-word, for example, 'flonty – xylophone', the individual is required to point to the real word in the pair. | $M = 53.72$<br>$SD = 5.28$<br>$Range = 24-60$ | Baddeley, Emslie & Nimmo-Smith, 1993 |
| | Sentence Comprehension (SntRec) | Listen to and judge grammatical acceptability of partial sentences, beginning with an (ambiguous, unambiguous) sentence stem (e.g., "Tom noticed that landing planes...") followed by a disambiguating continuation word (e.g., "are") in a different voice. Ambiguity is either semantic or syntactic, with empirically determined dominant and subordinate interpretations | $M = 0.89$<br>$SD = 0.07$<br>$Range = 0.46-1$ | Rodd, Longe, Randall, & Tyler, 2010 |
| | Picture-Picture Priming (PicNam) | Name the pictured object presented alone (baseline), then when preceded by a prime object that is phonologically related (one, two initial phonemes), semantically related (low, high relatedness), or unrelated | $M = 0.78$<br>$SD = 0.09$<br>$Range = 0.5-0.94$ | Clarke, Taylor, Devereux, Randall, & Tyler, 2013 |
| | Verbal Fluency (VrbFl) | Mean of letter (phonemic) fluency and animal (semantic) fluency task. For phonemic fluency task, participants have 1 min to generate as many words as possible beginning with the letter 'p'. For semantic fluency task, participants have 1 min to generate as many words as possible in the category 'animals'. | $M = 20.72$<br>$SD = 5.4$<br>$Range = 6-37.5$ | Lezak, Muriel, & Deutsch, 1995 |
| | Proverb Comprehension (ProV) | Read and interpret three English proverbs. | $M = 4.54$<br>$SD = 1.62$<br>$Range = 0-6$ | Hodges, 1994 |
| Emotional Processing | Face Recognition (FaceRec) | Given a target image of a face, identify same individual in an array of 6 face images (with possible changes in head orientation and lighting between target and same face in the test array) | $M = 22.93$<br>$SD = 2.32$<br>$Range = 14-27$ | Benton, 1994 |
| | Emotion Expression Recognition (EmoRec) | View face and label emotion expressed (happy, sad, anger, fear, disgust, surprise) where faces are morphs along axes between emotional expressions. | $M = 86.6$<br>$SD = 10.74$<br>$Range = 47.5-100$ | Ekman & Friesen, 1976 |
| Memory | Visual Short-Term Memory (VSTM) | View (1–4) coloured discs briefly presented on a computer screen, then after a delay, attempt to remember the colour of the disc that was at a cued location. | $M = 2.43$<br>$SD = 0.58$<br>$Range = 0-3.5$ | Zhang & Luck, 2008 |
| | Story Recall (StrRec) | Listen to a short story, recall freely immediately after, then again after a delay, and finally answer recognition memory questions. Delayed recall measure used here. | $M = 12.98$<br>$SD = 4.23$<br>$Range = 0-24$ | Wechsler, 1999 |
| Processing Speed | Choice Motor Speed (MRSp) | Time-pressured movement of a cursor to a target by moving an (occluded) stylus under veridical, perturbed (30°), and reset (veridical again) mappings between visual and real space. | $M = 0.19$<br>$SD = 0.06$<br>$Range = 0.05-0.85$ | |
| | Choice Motor Coefficient of Variation (MRCv) | Standard deviation divided by mean of reaction time of choice motor speed. Reflects the relative measure of variability. | $M = 1.84$<br>$SD = 0.38$<br>$Range = 0.91-2.98$ | |

### MEG data acquisition and pre-processing

Data were collected using a 306-channel VectorView MEG system (Elekta Neuromag, Helsinki), consisting of 102 magnetometers and 204 orthogonal planar gradiometers, located in a magnetically shielded room. MEG resting state data (sampled at 1 kHz with a highpass filter of 0.03 Hz) were recorded for 8 min and 40 s, while participants remained still in a seated position with their eyes closed. Head position within the MEG helmet was estimated continuously using four Head-Position Indicator (HPI) coils to allow offline correction of head motion. Two pairs of bipolar electrodes recorded vertical and horizontal electrooculogram (VEOG, HEOG) signals to monitor blinks and eye-movements, and one pair of bipolar electrodes recorded the electrocardiogram (ECG) signal to monitor pulse-related artefacts.

The MaxFilter 2.2.12 software (Elekta Neuromag Oy, Helsinki, Finland) was used to apply temporal signal space separation (tSSS, Taulu et al., 2005) to the continuous MEG data to remove noise from external sources (correlation threshold 0.98, 10-sec sliding window), to continuously correct for head-motion (in 200-ms time windows), to remove mains-frequency noise (50-Hz notch filter), and to detect and reconstruct noisy channels. Following these de-noising steps, data were imported into Matlab (The MathWorks, Inc.) and preprocessed using a mixture of SPM12 (http://www.fil.ion.ucl.ac.uk/spm) and the OHBA Software Library (OSL; https://ohba-analysis.github.io/osl-docs/). Bad segments were detected and rejected by identifying outliers in the standard deviation of the signal using the Generalized ESD test^46^ at a significance level of a 0.1 (mean % time points rejected = 1.44, *SD* = 1.35). Data were then down-sampled to 200Hz, and a band pass filter applied from 1–45 Hz to remove slow trends and high frequency noise.

### MEG source reconstruction, parcellation, and envelope calculation

The MEG data were co-registered to each participant’s structural T1-weighted MRI, using three anatomical fiducial points (nasion, and left and right pre-auricular points) that were digitized for the MEG data and identified manually on the MRIs. The median distance between each scalp headshape point and its nearest vertex was calculated for each participant, and those with a median distance greater than 6 mm (*n*=15; see *Participants* section above) were excluded from subsequent analyses.

Source space activity was then estimated for each participant at every point of an 8 mm whole-brain grid using a single-shell lead-field model and a linearly constrained minimum variance (LCMV) scalar beamformer^47,48^, combining data from both magnetometers and gradiometers. Source-reconstructed time-series (in each epoch) for each grid point were then parcelled into 38 regions of interest (ROIs; defined by selecting a subset of 19 of the ROIs in the Harvard–Oxford cortical brain atlas, available in FSL, and splitting each into two lateral halves to create 38 binary ROIs^49^) in order to reduce the dimensionality of the oscillatory activity submitted to the HMM (see below). Each parcel was represented by the first principal component across grid points within an ROI, and magnetic field spread between parcels was reduced by symmetric, multivariate orthogonalization^49,50^. Next, the amplitude envelope of each parcel’s time-course was calculated using a Hilbert transform, subsequently down-sampled to 20 Hz for computational efficiency.

### Group level exploratory analysis of networks (Hidden Markov Model)

The HMM procedure assumes that the same set of microstates underpin the HMM-derived states across participants. Therefore, demeaned and normalized envelope data for each participant were concatenated temporally across all participants to produce a single dataset, using the group-level exploratory analysis of networks (GLEAN) toolbox (https://github.com/OHBA-analysis/GLEAN^25^).

We set the analysis a-priori to derive 8 states from the data, based on previous work suggesting that this number represents a reasonable trade-off between a sufficiently rich but not overly complex representation^21^. HMMs describe the dynamics of brain activity as a sequence of transient events, each of which corresponds to a visit to a particular brain state. Each state describes the data as coming from a unique 38-dimensional multivariate normal distribution, defined by a covariance matrix and a mean vector. Therefore, each state corresponds to a unique pattern of amplitude envelope variance and covariance that reoccurs at different time points. The HMM state time-courses then define the points in time at which each state was “active” or “visited”. We obtained these estimated state time-courses, represented by a binary sequence showing the points in time when that state was most probable, using the Viterbi algorithm^51^. We then partially correlated the time-courses onto whole-brain parcel-wise amplitude envelopes concatenated across subjects in order to produce spatial maps of the changes in amplitude envelope activity associated with each state. The resulting state maps show the brain areas whose amplitude envelopes increase or decrease together (covary) when that state is active, compared to what happens on average over time.

Using the state time-courses, we quantified the temporal characteristics of each state according to four measures of interest: (1) Fractional Occupancy (FO): the proportion of time the state was active; (2) Mean Life Time (MLT): the average time spent in the state before transitioning to another state; (3) Number of Occurrences (NO): the number of times the state was active; and (4) Mean Interval Length (MIL): the average duration between recurring visits to that state.

### Relating HMM states to age and cognition (Canonical Correlation Analyses)

To identify how the temporal characteristics of the HMM states relate to age and cognition, we used Canonical Correlation Analyses (CCA^52–54^; [see Figure 1 in Wang et al^54^ for schematic illustration of CCA]). CCA is a multivariate technique that can identify and measure linear relations between two sets of variables. Linear combinations within each of the sets are defined such that the correlations of these combinations between sets (e.g., between HMM profile, and cognitive profile) are maximized, resulting in CCA components, or “modes”. This multivariate approach is useful when the observed variables within each set are correlated (as is the case for the above HMM temporal characteristics, and for the cognitive scores).

CCA was employed via *canoncorr* in Matlab. The number of modes produced by this analysis is always equal to the number of variables in the small dataset (though not all modes necessarily explain a substantial portion of the variance, see *Results* section above). In canoncorr’s terminology (see also studies mentioned above^52–54^), each CCA mode is associated with canonical “coefficients” across the variables in each set and “scores” across the observations (participants). This correlation between the participant scores of each set (for a given mode) is termed the “canonical correlation” (denoted by *R_c_*), and its squared value (*R^2^_c_*) represents the proportion of variance shared between the variable sets. The correlation between the participant scores and each original variable (for a given set and given mode) is called the “structure coefficient” (denoted by *r_s_*), and the set of structure coefficients represents the “profile” of the CCA mode across those variables. Structure coefficients are often used to guide interpretation of multivariate analyses, and are particularly useful in the presence of multicollinearity^28^.

All variables were z-scored before subjected to CCA, in order to make the various parameters more comparable across variables. First, we used CCA to relate the 4 HMM measures across all 8 states (Set 1, 32 variables) to age. Note that in this case, the CCA analysis is equivalent to multiple linear regression (MLR) because the second set contains a single variable (age). Nevertheless, for consistency with subsequent analysis, we used CCA rather than MLR. We then conducted another CCA analysis to relate the 32 HMM measures (Set 1) to the 13 cognitive measures (Set 2). Having used CCA to establish relationships between the HMM brain measures and the cognitive measures across all ages, we then asked whether the relationship between HMM profile and cognitive profile (i.e., the relations between canonical scores obtained by the final CCA) varied with age, using a moderation analysis (see previous studies^39,55^ for a similar approach with different measures). Specifically, we constructed a multiple linear model where HMM scores (for a given mode), age and their interaction term (HMM profile × age) were used as independent variables and cognitive scores (for that mode) as the dependent variable (all statistical tests were two-sided). In order to visualise the results of this continuous moderation analysis, we created scatter plots of HMM profile versus cognitive profile for six equally-sized age groups (*n* = 99 in each group).

### Additional control analyses

We performed several additional analyses in order to ensure that the results are robust and interpretable. First, some of the temporal measures of the HMM states (and MIL in particular) included a relatively large number of outliers (see Figure 3). Therefore, in order to ascertain that the results are not biased by these outliers, we repeated the main CCA and moderation analyses after excluding 98 participants depicting outliers (*SD*>3, n=496) in one or more measures. Following this removal, the first CCA mode remained highly significant (*R_c_* = .512, *R^2^_c_* = 26%, *p* < .001), and the pattern of structure correlations remained remarkably similar to that observed with the full sample (see Supplementary Figure 2). The results of the moderation analysis were also similar, with a significant association between HMM and cognitive profiles after accounting for the main effect of age (*β* = .16, *t*(492) = 5.18, *p* < .001), together with a significant interaction between age and HMM profile (*β* = .13, *t*(492) = 4.84, *p* < .001).

Moreover, in order to confirm the significance of *R_c_* of the first CCA mode as determined by parametric assumptions, we also estimated *R_c_* against a distribution of 10,000 correlation coefficients based on permuting across participants their canonical scores for the cognitive data. The canonical coefficient for the true data (*R_c_* = .49) was greater than for any of the random permutations (equivalent to *p* < .0001, see Supplementary Figure 3). We also carried out a separate cross-validation analysis in which the CCA was only run on a subset of the data, and the outputs tested against the rest. In each of 10,000 iterations, we randomly chose 80% of the subjects for the “training” subset, leaving the other 20% for the “testing” subset. The canonical correlations across training sets for the first mode was similar to the original result (ranging between .45 and .56). For each iteration, we took the CCA HMM and cognition weight vectors for the first mode from the training subset, and multiplied them by the testing subset, in order to estimate participant scores for HMM and cognition. We then computed the correlation between these scores to estimate the canonical coefficient for the testing subset. Mean *R_c_* was .32 (mean *p* = .007, see Supplementary Figure 3 for the full distribution). Importantly, the canonical coefficient was significant at *p* < .05 for 96.7% of the testing subsets and at p < .001 for 63.7% of the testing subsets. Taken together, the results of these analyses confirm that the first CCA mode was highly significant.

Finally, we repeated the HMM-age CCA analysis with an additional quadratic term, in order to account for potential non-linear (quadratic) age-effects. We computed a quadratic age term, orthogonal to the linear age term, and ran a CCA analysis that relates the 4 HMM measures across all 8 states (Set 1, 32 variables) to both age terms (Set 2, 2 variables). Both possible CCA modes were significant (*p* < .001) and explained 28.6% and 12.3% of the shared variance, respectively. The structure coefficients (r_s_) are shown in Table 3 below. As apparent from the table, the first CCA mode captured the linear relations between the HMM measures and age, whereas the second CCA mode captured the quadratic relations. Importantly, we have observed an age-related shift from lower to higher-order states (captured by the first CCA mode), which was nearly identical to that observed in our initial analysis (which did not include the quadratic age term). This suggests that this neural profile remains stable even when accounting for non-linear age effects.

**Table 3.** Structure coefficients for the CCA relating HMM measures with linear and quadratic age (N=594)

| CCA Mode I |  |  |  |  |  |  |
| --- | --- | --- | --- | --- | --- | --- |
| State | FO | MLT | NO | MIL | linAge | quadAge |
| FTP1 | .27** | .14** | .28** | -.12** | .99** | .02 |
| FTP2 | .23** | .15** | .24** | -.21** |  |  |
| FTP3 | .19** | .44** | .11* | -.07 |  |  |
| HOV | .24** | .16** | .35** | -.08* |  |  |
| EV1 | -.35** | -.51** | -.00 | -.01 |  |  |
| EV2 | -.45** | -.38** | -.33** | .18** |  |  |
| SM1 | .18** | .04 | .22** | -.13** |  |  |
| SM2 | -.03 | -.08* | .00 | .02 |  |  |
| CCA Mode II |  |  |  |  |  |  |
| State | FO | MLT | NO | MIL | linAge | quadAge |
| FTP1 | -.04 | -.35** | .05 | .01 | -.02 | .99** |
| FTP2 | .04 | -.15** | .1* | -.05 |  |  |
| FTP3 | .52** | .25** | .46** | -.46** |  |  |
| HOV | -.42** | -.5** | -.31** | .25** |  |  |
| EV1 | .21** | .07 | .17** | -.27** |  |  |
| EV2 | .08 | -.11* | .19** | -.19** |  |  |
| SM1 | .1* | -.32** | .21** | -.3** |  |  |
| SM2 | -.04 | -.27** | .06 | -.11* |  |  |

\* $p < .05$ , \*\* $p < .005$ . Note: Structure coefficients ( $r_s$ ) for two CCA modes are presented. Coefficients are shown for each of the 4 HMM measures (FO, MLT, NO, MIL), for each state.

## Data Availability Statement

Raw data from the Cam-CAN project is available from the managed-access portal at http://camcan-archive.mrc-cbu.cam.ac.uk, subject to conditions (see website). For a complete description of Cam-CAN data and pipelines, see Shafto et al.^40^ and Taylor et al.^56^. In addition, pre-processed mean data used for analyses and figures will be available on OSF.

## Code Availability Statement

All code used for the analyses reported in the manuscript is available for peer-review on OSF, and will be made publically available upon acceptance of the manuscript.

## Acknowledgments

The Cambridge Centre for Ageing and Neuroscience (Cam-CAN) research was supported by the Biotechnology and Biological Sciences Research Council (BB/H008217/1); R.T is supported by a British Academy Postdoctoral Fellowship (SUAI/028 RG94188); K.A.T is supported by British Academy (PF160048) and the Guarantors of Brain (G101149); R.H is supported by a UK Medical Research Council grant (SUAG/010 RG91365). We are grateful to the Cam-CAN respondents and their primary care teams in Cambridge for their participation in this study. We also thank colleagues at the MRC Cognition and Brain Sciences Unit MEG and MRI facilities for their assistance. The Cam-CAN corporate author consists of the project principal personnel: Lorraine K Tyler, Carol Brayne, Edward T Bullmore, Andrew C Calder, Rhodri Cusack, Tim Dalgleish, John Duncan, Richard N Henson, Fiona E Matthews, William D Marslen-Wilson, James B Rowe, Meredith A Shafto; Research Associates: Karen Campbell, Teresa Cheung, Simon Davis, Linda Geerligs, Rogier Kievit, Anna McCarrey, Abdur Mustafa, Darren Price, David Samu, Jason R Taylor, Matthias Treder, Kamen Tsvetanov, Janna van Belle, Nitin Williams; Research Assistants: Lauren Bates, Tina Emery, Sharon Erzinc, jioglu, Andrew Gadie, Sofia Gerbase, Stanimira Georgieva, Claire Hanley, Beth Parkin, David Troy; Research Interviewers: Jodie Allen, Gillian Amery, Liana Amunts, Anne Barcroft, Amanda Castle, Cheryl Dias, Jonathan Dowrick, Melissa Fair, Hayley Fisher, Anna Goulding, Adarsh Grewal, Geoff Hale, Andrew Hilton, Frances Johnson, Patricia Johnston, Thea Kavanagh-Williamson, Magdalena Kwasniewska, Alison McMinn, Kim Norman, Jessica Penrose, Fiona Roby, Diane Rowland, John Sargeant, Maggie Squire, Beth Stevens, Aldabra Stoddart, Cheryl Stone, Tracy Thompson, Ozlem Yazlik; and administrative staff: Dan Barnes, Marie Dixon, Jaya Hillman, Joanne Mitchell, Laura Villis.

## Author contribution

R.T: directed the project together with R.H; designed, planned, and carried out the analyses; wrote the paper with input from all authors. K.A.T: consulted on CCA analyses; provided input on the paper; D.P: consulted on pre-processing; provided input on the paper; D.N; provided input on results at early stages; provided input on the paper. R.H: directed the project together with R.T; commented on structure and analysis approach; provided input on the paper.

## Supplementary Information

**Supplementary Figure 1.**
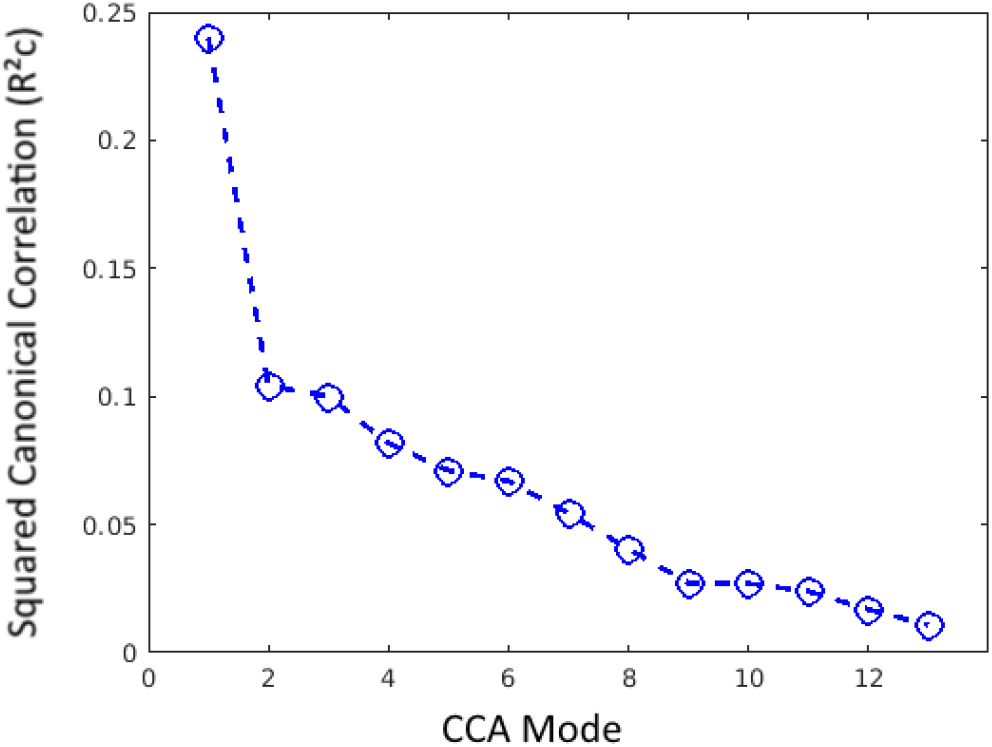
Squared canonical correlation (*R^2^c*) for each CCA mode, which represents the proportion of variance shared by the two synthetic variates. Only the first CCA mode, which accounted for a substantial portion of the shared variance (*R^2^c* = 24%), was interpreted.

**Supplementary Figure 2.**
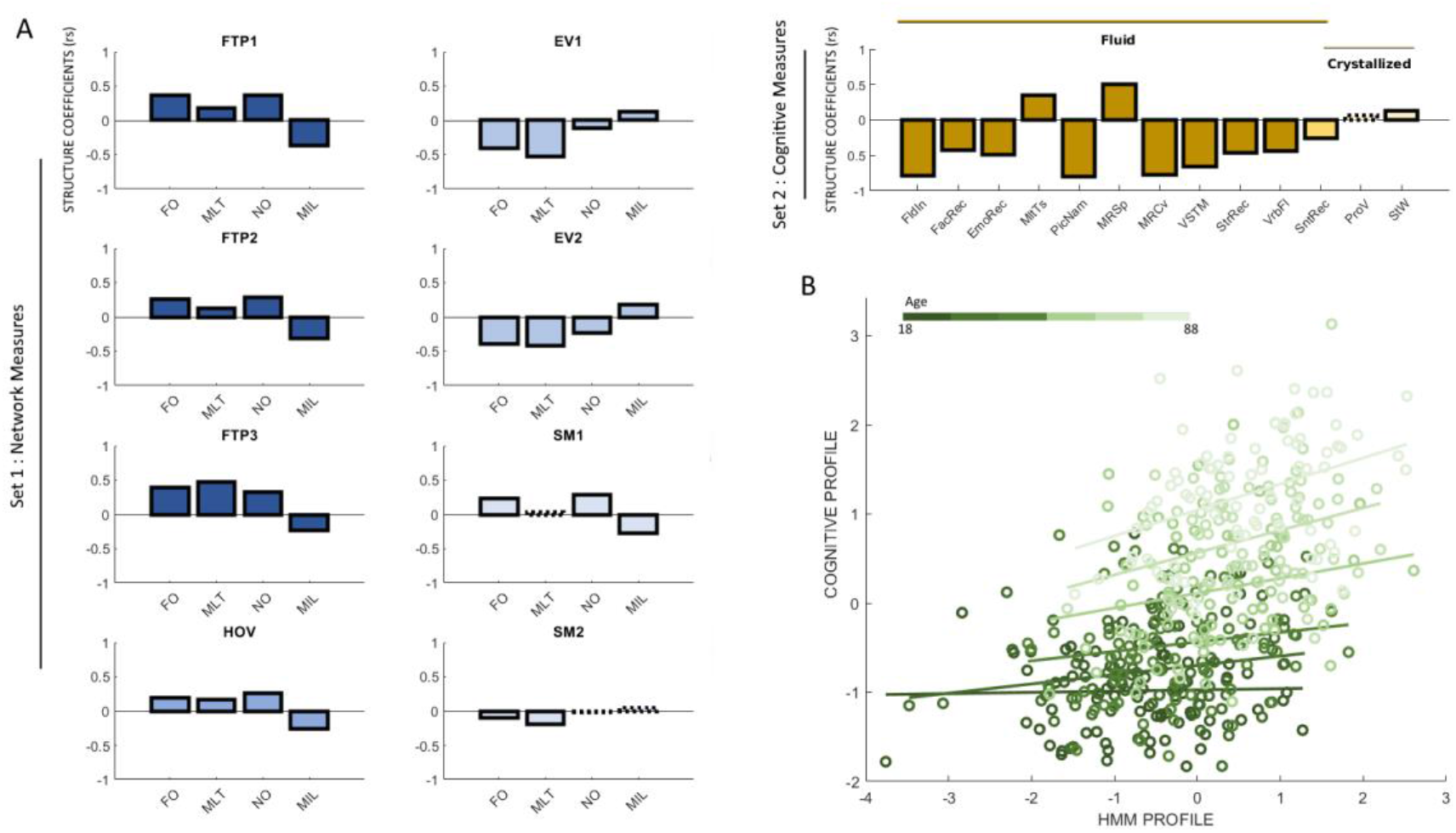
Outcomes of CCA and moderation analyses with n=496, following the removal of outliers (*SD* > 3 in one or more HMM measures).

**Supplementary Figure 3.**
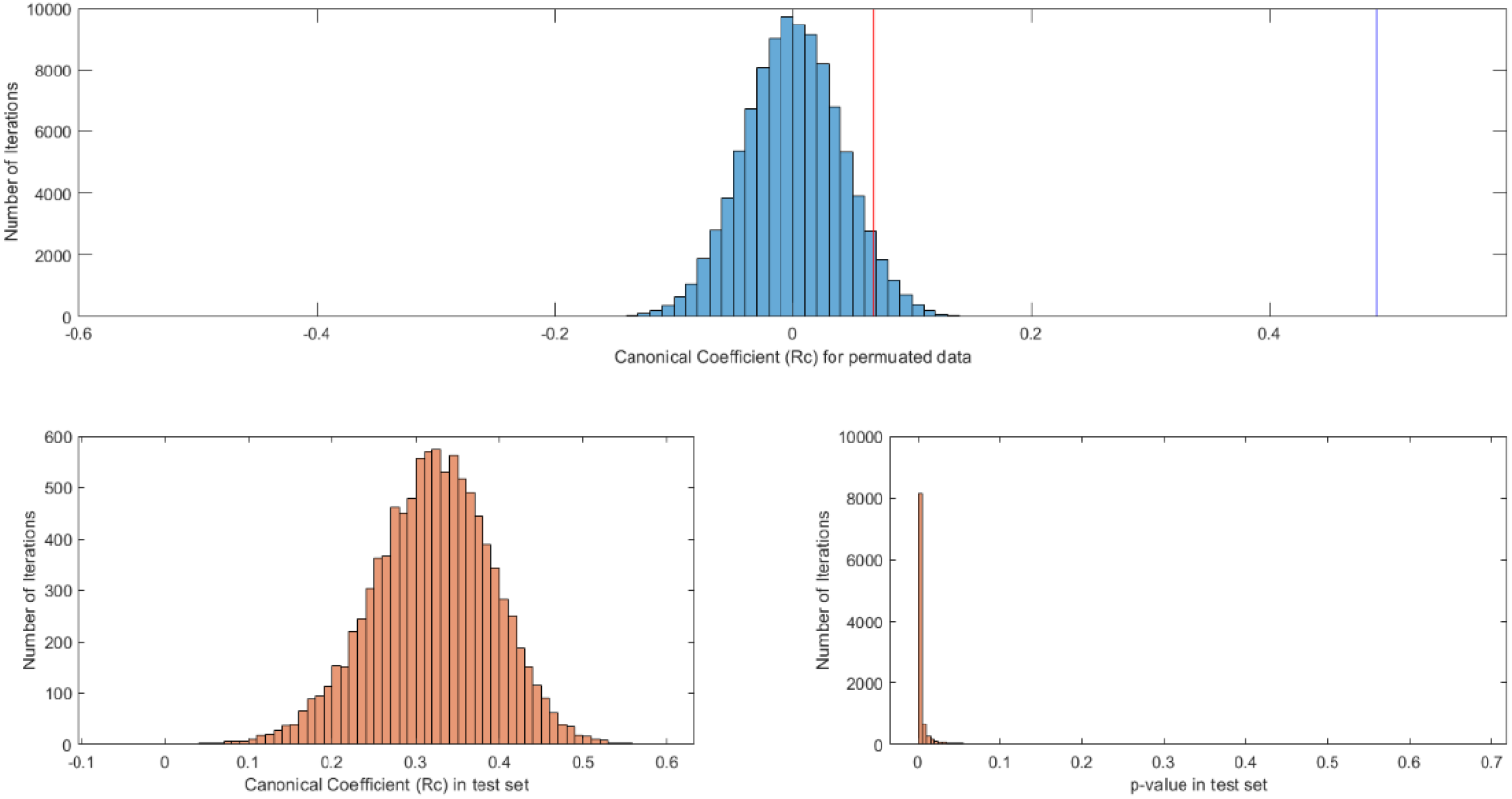
Outcomes of control analyses. Top: histogram of canonical coefficients (*R_c_*) of the first CCA mode for permutated data across 10,000 iterations. Red line indicates *R_c_* at 95% of the distribution of permutation data (equivalent to p = .05). Blue line indicates *R_c_* obtained with the actual data. Bottom: Histogram of Rc (left) and p-values (right) of the first CCA mode in the test set after applying weight vectors from the train set, across 10,000 iterations.

